# Microphysiological uremia model reveals biophysical potentiators of vascular dysfunction

**DOI:** 10.1101/2024.12.18.629161

**Authors:** Mitesh Rathod, Stephanie A. Huang, Wen Yih Aw, Elizabeth L. Doherty, Sara M. Meehan, Prabir Roy-Chaudhury, William J. Polacheck

## Abstract

Cardiovascular disease is a leading cause of mortality in individuals with chronic kidney disease. Hypertension, common among patients with chronic kidney disease, is a major contributor to both kidney damage and the heightened cardiovascular risk in these patients. Advanced chronic kidney disease is associated with elevated levels of circulating uremic toxins, particularly indoxyl sulfate and p-cresyl sulfate, and are known to exacerbate cardiovascular risk by promoting inflammatory processes, including monocyte adhesion, rolling, and extravasation. However, despite the established link between chronic kidney disease and cardiovascular disease, the specific role of uremic toxins in monocyte-endothelial interactions in hypertensive settings remains largely underexplored. In this study, we developed a 3D microfluidic model to examine the effects of indoxyl sulfate on monocyte adhesion and extravasation across engineered microvessels embedded in collagen hydrogels with different densities under controlled luminal pressure. We found that elevated pressure alone significantly enhanced monocyte adhesion and extravasation, regardless of matrix density, and that the uremic environment further increased these effects. Additionally, denser hydrogels primed THP-1 monocyte cells toward a pro-inflammatory like phenotype with reduced phagocytic capacity, while softer hydrogels induced an anti-inflammatory like phenotype with enhanced phagocytosis. However, exposure to the uremic environment diminished phagocytosis and shifted cells toward a pro-inflammatory like state, irrespective of matrix density. The presented approach has the potential to experimentally dissect multiple factors that contribute to elevated cardiovascular risks in chronic kidney disease patients and improve the understanding of mechanisms involved in monocyte dynamics in chronic kidney disease- related cardiovascular disease.

## 1. Introduction

Chronic kidney disease (CKD) impacts approximately 13.4% of the global population [1] and is a progressive condition that can eventually result in permanent kidney failure, requiring patients to undergo dialysis or receive a kidney transplant for survival. The number of individuals on dialysis is rapidly increasing worldwide, primarily due to an aging population and rising rates of hypertension and type 2 diabetes mellitus, which are significant contributors to the development and progression of CKD [2]. Patients with CKD face elevated cardiovascular risks that correlate with glomerular filtration rate (GFR): 75.3% of CKD patients in stages G4-G5 (GFR < 29 ml/min/1.73m^2^), 66.6% of CKD patients in stage G3 (30-59 ml/min/1.73m^2^), and 63.4% of CKD patients in stages G1-G2 (GFR ≥ 60 ml/min/1.73m^2^) experiencing cardiovascular disease (CVD), compared to 37.5% of those without CKD [3]. The elevated CVD risk is underscored by the common occurrence of atherosclerosis and the increased rates of peripheral artery disease, stroke, arrhythmias, and myocardial infarctions, leading to higher mortality rates from CVD [4–8].

Hypertension is a major global health issue and remains one of the top preventable causes of death [9, 10], It is the most frequent comorbidity of CKD, afflicting 67% - 92% of CKD patients, and becoming more prevalent and severe as kidney function declines [11]. The relationship between hypertension and CKD is complex, as hypertension can both result from and contribute to the progression of kidney disease. [12–14]. Irrespective of the underlying cause of CKD, an increase in systemic blood pressure accelerates the decline in glomerular filtration rate (GFR) [15, 16], establishing hypertension as an independent risk factor for end-stage renal disease (ESRD) [17, 18]. Both clinical and experimental evidence supports the notion that blood pressure is closely linked to kidney function; for instance, normotensive kidney recipients from hypertensive donors often develop hypertension in humans [19] and rats [20], while hypertensive recipients of normotensive kidneys may normalize their blood pressure in both human [21] and rats [22]. Moreover, hypertension not only accelerates kidney disease progression but also contributes to the onset and intensification of CVD. Previous studies have indicated that hypertension raises the likelihood of new or recurring cardiovascular events in individuals with stages 2–3 CKD [23].

One contributing factor to high blood pressure in CKD patients is alterations to the blood vasculature. Continuous exposure to dysregulated circulating factors including hormones and uremic toxins leads to vascular dysfunction, resulting in elevated blood pressure that is often resistant to medication [24]. Vascular dysfunction can occur early in CKD progression and while seeming to act independently from hypertension, plays a significant role in the subsequent development of high blood pressure [25–27]. For example, in CKD rat models, vascular changes can be structural, as characterized by vessel remodeling that enhances contractility and reduces elasticity of arteries, thus increasing vascular resistance [28–30]; or functional, such as impaired acetylcholine-induced relaxation [25, 27, 31, 32]. Moreover, arterial stiffness is heightened in CKD patients [33–36] and is linked to an independent rise in cardiovascular risk [35,36]. Interestingly, in patients with advanced CKD, a decline in aortic stiffness has been associated with improved survival, independent of blood pressure [35]. These vascular alterations are frequently linked to an overexpression of intercellular adhesion molecules like ICAM-1 and VCAM-1 [27, 37, 38]. Furthermore, these vascular modifications are associated with the development of cardiac pathology, in turn raising blood pressure and worsening as CKD advances [39]. Notably, endothelial dysfunction in CKD appears to stem from increased endothelial injury and diminished repair mechanisms [40], which are linked to various systemic blood pressure and hemodynamic changes.

In patients with advanced CKD, elevated serum levels of several circulating uremic toxins, especially indoxyl sulfate (IS) and p-cresyl sulfate (pCS), are observed alongside inflammatory markers [41]. These toxins are challenging to eliminate through standard dialysis procedures due to their strong binding affinity to blood proteins [42], leading to their accumulation in the bloodstream of those with advanced CKD [43]. There is considerable evidence indicating that uremic toxins predispose CKD patients to CVD by promoting monocyte adhesion, rolling, and extravasation. Intermediate monocyte subtypes in CKD exhibit a proatherogenic pattern, expressing chemokines and adhesion molecules that enhance adhesion to endothelial cells [44] IS has been shown to increase the adhesion of THP-1 monocytes to activated human endothelial cells [45], while pCS promotes oxidative burst activity in monocytes [46]. In cultures of endothelial cells and macrophages, pCS drives the expression of inflammatory factors and adhesion molecules through the production of reactive oxygen species (ROS), which has been observed in vivo during leukocyte-endothelium interactions [47]. Additionally, studies using intravital microscopy and rodent models have demonstrated that exposure to high concentrations of IS increases leukocyte rolling along endothelial surfaces and promotes leukocyte adherence and extravasation [45]. Collectively, these findings suggest that uremic toxins may facilitate monocyte migration and inflammation-related CVD in CKD. However, little research has been conducted on the processes of monocyte adhesion and migration in the context of CVD associated with CKD, and 3D in vitro models that recapitulate the hypertensive environment under uremic conditions are lacking.

To address these challenges, we introduce a 3D microfluidic system designed to study the impact of a uremic environment on inflammatory processes in the endothelium, with a specific focus on monocyte recruitment under hypertensive conditions. First, we assess how the poroelastic properties of collagen type I hydrogels influence monocyte adhesion in response to luminal pressure. Building on this analysis, we investigate the effects of indoxyl sulfate on monocyte adhesion and extravasation across microvessels embedded in collagen with varying densities under elevated luminal pressure and identify the adhesion receptors involved in monocyte recruitment at the endothelium. Finally, we explore how the properties of the collagen hydrogel and uremic environment modulate the phenotype and phagocytic capacity of THP-1 monocytes.

## 2. Results

### 2.1 Microfluidic model for modulating luminal and transmural pressure

Arterial stiffness and hypertension are both increased in patients with CKD [48]. To investigate the role of ECM stiffness and luminal pressure on monocyte adhesion and invasion in the context of CKD, we microfabricated microfluidic devices with two parallel microchannels embedded in 2.5 and 6 mg/mL collagen type I hydrogels (**Figure 1A**). We then cultured THP-1 monocytes in microchannels under luminal pressures of 0, 20 and 200 Pa (**Figure 1B-C**). For both 2.5 mg/ml and 6 mg/ml collagen concentrations, the number of adhered monocytes to the microchannel wall increased with increasing luminal pressure (**Figure 1B-C**), and at all pressures, the number of adhered monocytes decreased in 6 mg/ml compared to 2.5 mg/mL collagen (**Figure 1C**). We hypothesized that the reduced monocyte adhesion to 6 mg/mL collagen hydrogels was due to differences in transmural flow and resultant drag on monocytes toward the wall of the channel . To characterize the poroelastic properties of the microvessels, we attached fluid reservoirs to apply a 200 Pa hydrostatic pressure head to one channel, while keeping the other channel at a constant atmospheric pressure (**Figure 1D-E, Figure S1**). The difference in column height of the source and sink channels was measured every 30 minutes for 14 hours (**Figure 1D**), and by fitting the pressure drop vs. time data to Darcy’s law (Supplementary Information), we calculated hydraulic conductivity of 2.5 mg/mL and of 6 mg/mL collagen hydrogels to be 1.46 × 10^-10^ m^2^/Pa s and 6.10 × 10^-11^ m^2^/Pa s, respectively (**Figure 1E**), consistent with previous measurements in other systems [49, 50].

**Figure 1.**
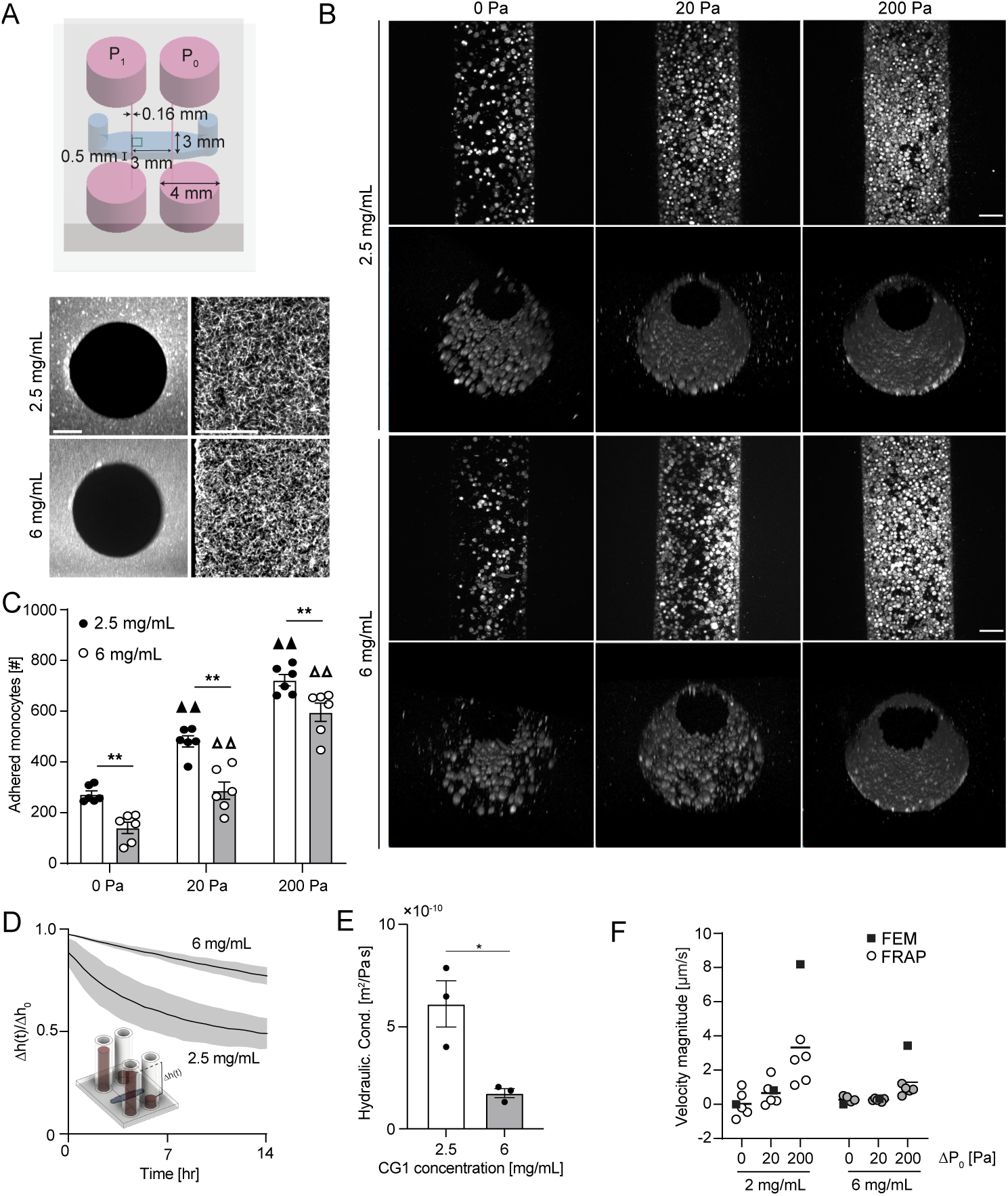
- Porous media microfluidic device for modulating luminal pressure. **A**) Schematic of 2-channel device with critical dimensions labeled and images of periluminal collagen hydrogel at two different mass densities (grey – AF647-labeled collagen type I, scale bar 25 μm). **B)** Monocyte adhesion and invasion into collagen hydrogels as determined by confocal imaging. Maximum intensity projections and 3D reconstruction of THP-1 monocytes in 2.5 mg/ml and 6 mg/ml collagen microchannels subjected to 0, 20 and 200 Pa luminal pressure (grey – phalloidin, scale bar 50 µm)**. C)** Quantification of adhered THP-1 monocyte to 2.5 mg/ml and 6 mg/ml collagen microchannel walls for different pressure conditions (3 microvessels per condition; ▲▲ p < 0.01 versus 0 Pa pressure for 2.5 mg/ml collagen, Δ Δ p < 0.01 versus 0 Pa pressure for 6mg/ml collagen, **p < 0.01, two-way ANOVA with a Bonferroni post-hoc test). **D**) Plot of normalized change in height between source and sink reservoir across collagen hydrogel used to calculate **E**) the hydraulic conductivity of collagen hydrogels within microfluidic devices. **F**) Interstitial fluid velocity magnitude as determined by FRAP of 70 kDa fluorescent dextran compared to expected values from FEM models (all bar graphs are mean ± SEM).

Using these values, we developed a computational fluid dynamics model using a Brinkman medium constitutive law in COMSOL to determine the fluid velocity field within the collagen hydrogel for applied pressure at the medium ports of the devices (**Figure S2**), and we compared the local velocity vector magnitude at the center of the hydrogel to that measured experimentally by FRAP of 70 kDa fluorescent dextran. For 20 and 200 Pa, we observed reduced transmural fluid velocity magnitudes in 6 mg/mL collagen hydrogels as compared to 2.5 mg/mL, consistent with the hypothesis that differences in flow magnitude underlie differences in monocyte adhesion, while at 200 Pa, there was a divergence in the expected and measured velocity magnitudes that was attenuated with increased collagen concentration (**Figure 1F**), consistent with our previous observations that the hydraulic conductivity of 2.5 mg/mL decreases with increasing pressure [51]. Previous work in poroelastic media has identified that the hydraulic conductivity of tissue is dependent on the applied pressure via changes in microstructure induced by fluid drag on the solid-phase ECM [52], and we observed that strain in ECM fibrils was dependent on transmural flow magnitude (**Figure S3**). Collectively, these results suggest that the poroelastic properties of the collagen hydrogel regulate monocyte adhesion in response to luminal pressure.

### 2.2 Monocyte adhesion and transmigration across the microvascular wall

Previously, we observed that in the absence of transmural pressure, engineered microvessels were more permeable to diffusion of 70 kDa dextran when cultured in 6 mg/mL collagen hydrogels as compared to 2.5 mg/mL, and that monocyte extravasation correlated with microvascular permeability [51]. To evaluate the effect of transmural pressure and the underlying extracellular matrix poroelastic properties on monocyte adhesion and transmigration across the microvascular wall, we seeded microvessels with HUVECs prior to the introduction of THP-1 monocytes (**Figure 2A, B**). Luminal pressures of 20 Pa and 200 Pa significantly increased the number of attached monocytes to the microvessel wall when compared to 0 Pa for both 2.5 mg/ml and 6 mg/ml collagen hydrogels (**Figure 2B**). The number of attached monocytes did not vary significantly as a function of collagen concentration for 0 and 20 Pa luminal pressures. However, at 200 Pa luminal pressure, we observed an increase in monocyte adhesion in 6 mg/mL as compared to 2.5 mg/mL collagen hydrogels (**Figure 2B**), which was opposite to the trend for monocyte culture without HUVECs, suggesting an active mechanism mediating monocyte adhesion to the endothelium. Interestingly, this increase in monocyte adhesion did not correlate with microvessel permeability to 70 kDa dextran, which remained higher in 2.5 mg/mL as compared to 6 mg/mL collagen hydrogels under applied pressure (**Figure S4**).

**Figure 2.**
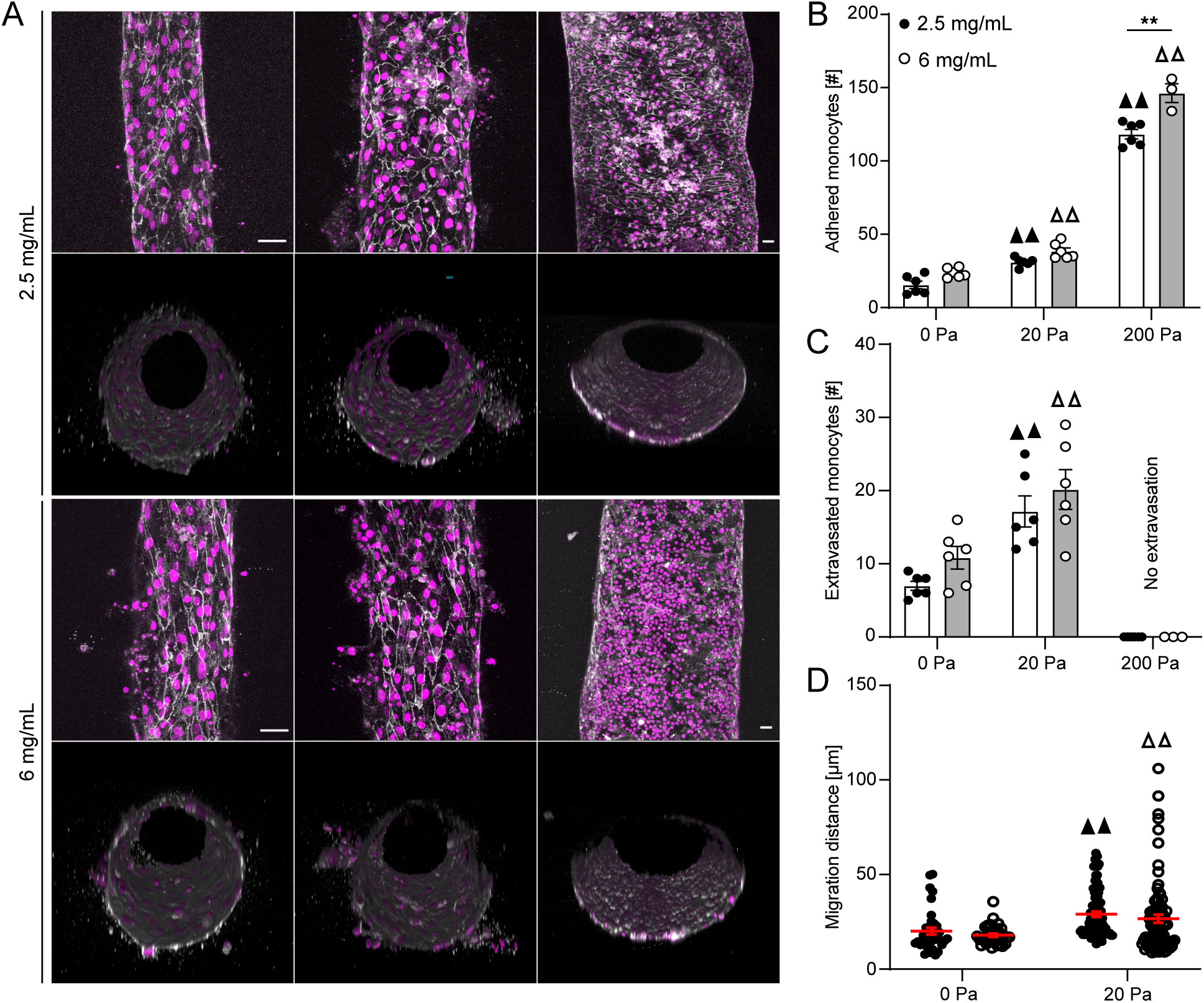
- Monocyte adhesion and transmigration across microvascular wall. **A**) 3D reconstruction confocal micrographs of HUVEC microvessels stained for DAPI (magenta) and VE-cadherin (white) and THP-1 monocytes stained for DAPI (magenta). Quantification of **B**) adhered monocytes to HUVEC microvessel, **C**) extravasated monocytes across HUVEC microvascular wall, and **D**) migration distance after extravasation in 2.5 mg/ml and 6 mg/ml collagen hydrogels subjected to 0, 20 and 200 Pa luminal pressure (three microvessel per condition; ▲▲ p < 0.01 versus 0 Pa pressure gradient for 2.5 mg/ml collagen, Δ Δ p < 0.01 versus 0 Pa pressure gradient for 6 mg/ml collagen, two-way ANOVA with a Bonferroni post-hoc test. All plots, mean ± SEM. Scale bars 50 µm).

Matrix stiffness has been shown to affect the migration patterns of THP-1 monocytes [53].We thus quantified the number of monocytes that extravasated from the microvessel into the surrounding ECM, and interestingly, 20 Pa luminal pressure significantly increased the monocyte extravasation across the microvessel wall for both collagen concentrations as compared to 0 Pa luminal pressure, while no transmigration was observed at 200 Pa for either concentration (**Figure 2C**). Furthermore, 20 Pa luminal pressure resulted in significantly longer invasion distances into the surrounding matrix as compared to 0Pa pressure gradient for both collagen concentrations (**Figure 2D**). Irrespective of the applied pressure gradient, the distance migrated by the THP-1 monocyte was overall higher in 2.5 mg/mL as compared to 6 mg/mL collagen, suggesting differential mechanisms driving monocyte adhesion and migration.

### 2.3 Monocyte adhesion and transmigration across the endothelium in uremic environment

To simulate the CKD microenvironment, we repeated THP-1 perfusion through bare collagen hydrogels in the presence of 2.5 µg/mL indoxyl sulfate, mimicking CKD stage 3 [54], and we observed a similar pressure-dependent increase in monocyte adhesion, and a reduction in adhesion in 6 mg/mL vs. 2.5 mg/mL collagen hydrogels. (**Figure 3A-C)**. Interestingly, indoxyl sulfate decreased monocyte adhesion in the absence of luminal pressure, with a 3.5x reduction at 0 Pa and 2.5 mg/mL collagen and a 4.5x reduction at 0 Pa and 6 mg/mL collagen (**Figures 1C, 3C**). However, in the presence of ECs, luminal pressures of 20 Pa and 200 Pa significantly increased monocyte attachment to the microvessel wall and higher collagen concentration resulted in greater monocyte attachment (**Figure 3D-F**), suggesting that the endothelium plays an active role in mediating monocyte attachment in the presence of IS and that there is a synergistic effect of luminal pressure and IS on monocyte attachment to the microvessel wall.

**Figure 3.**
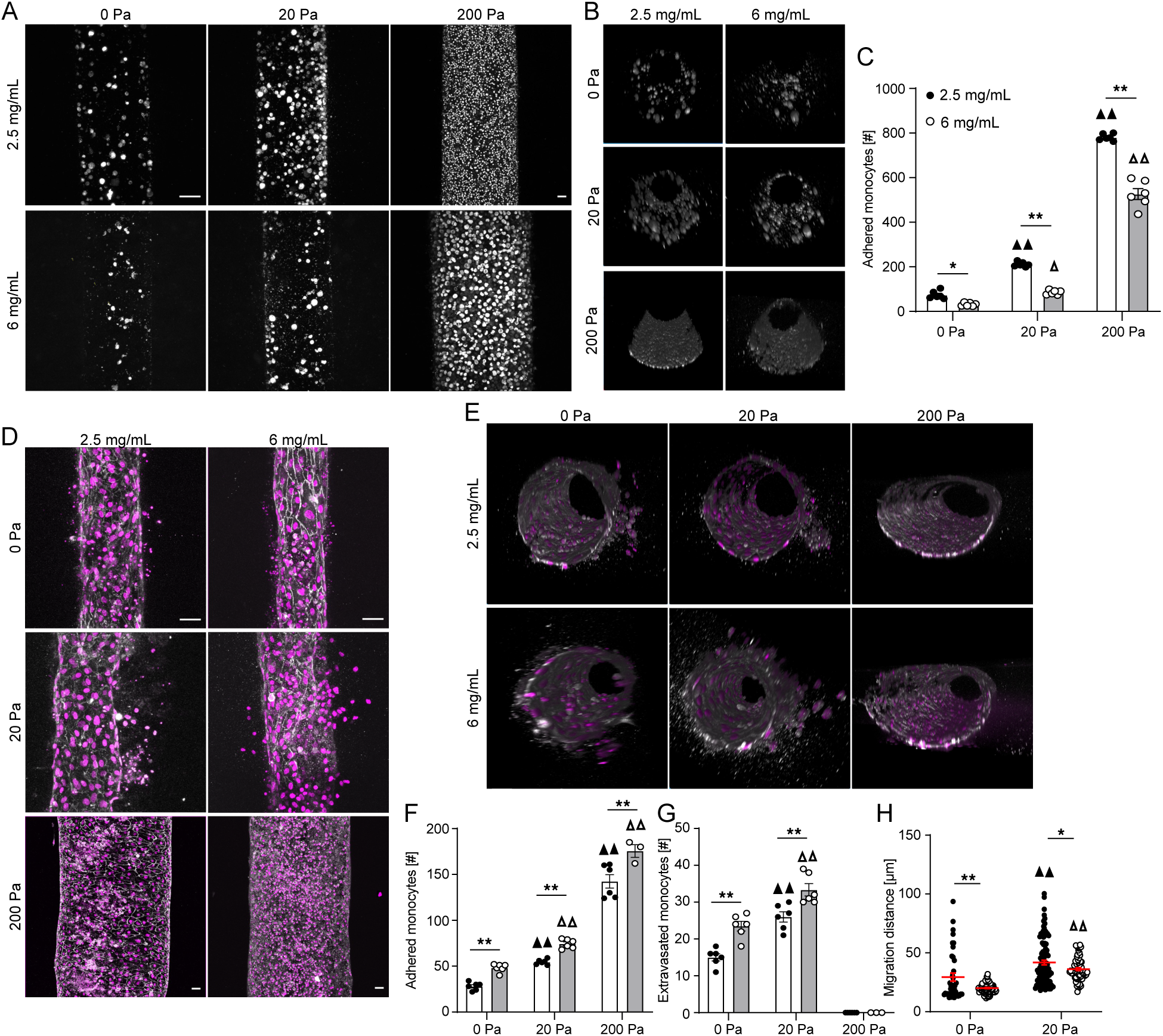
- **Monocyte adhesion and invasion into collagen hydrogels in a uremic microenvironment. A**) Confocal maximum intensity projections and **B)** 3D reconstructions of THP-1 monocytes cultured in 2.5 mg/ml and 6 mg/ml collagen microchannels subjected to 0, 20 and 200 Pa pressure conditions and treated with 2.5 µg/ml indoxyl sulfate (grey – phalloidin). **C)** Quantification of the number of monocytes adhered to the microvessel wall. **D)** Confocal maximum intensity projections and **E)** 3D reconstructions of HUVECs stained for DAPI (magenta) and VE-cadherin (white) and THP-1 monocytes stained for DAPI (magenta). Quantification of **F)** adhered monocytes to endothelium, **G)** extravasated monocytes across endothelial wall, and **H)** monocyte migration distance in 2.5 mg/ml and 6 mg/ml collagen hydrogels in response to 2.5 µg/ml indoxyl sulfate treatment under various pressure conditions (three microvessel per condition; ▲▲ p < 0.01 versus 0 Pa pressure for 2.5 mg/ml collagen, Δ p < 0.05, Δ Δ p < 0.01 versus 0 Pa pressure for 6 mg/ml collagen, *p < 0.05, **p < 0.01, two-way ANOVA with a Bonferroni post-hoc test. All plots, mean ± SEM. Scale bar 50 µm).

Treatment with IS also resulted in a significant increase in monocyte extravasation for both collagen densities for 20 Pa luminal pressure compared to 0 Pa (**Figure 3G**). Moreover, the extravasation was significantly higher in 6 mg/mL hydrogels compared to 2.5 mg/mL hydrogels at lower magnitude luminal pressure, while application of 200 Pa luminal pressure resulted in no extravasation irrespective of the collagen concentration (**Figure 3G**). Interestingly, treatment of the microvessel and monocytes with IS resulted in overall higher numbers of attached monocytes and extravasation when compared to the untreated conditions (**Figures 2B-C, 3F-G**). IS treatment also resulted in significantly longer migration distances for both collagen concentrations at 20 Pa luminal pressure compared to 0 Pa (**Figure 3H**). Moreover, in a similar trend as the untreated condition (**Figure 2D**), the 2.5 mg/mL collagen hydrogels resulted in significantly higher extravasation distances when compared to 6 mg/mL for both the luminal pressure values (**Figure 3H**), suggesting that the mechanisms regulating monocyte adhesion and subsequent migration are independent.

### 2.4 Effects of IS on expression of adhesion receptors by the endothelium

Motivated by the results that demonstrate ECs promote active recruitment of monocytes in the presence of IS, we assayed for cell adhesion markers previously shown to modulate interactions between monocytes and ECs. We measured protein expression via western blot for HUVECs cultured on 2.5 mg/mL and 6 mg/mL collagen hydrogels with and without IS treatment (**Figure 4A**). We observed a significant increase in ICAM-1 expression as a function of both collagen density and IS treatment, consistent with monocyte adhesion results (**Figure 4B**). VCAM1 expression was significantly higher on 6 mg/mL collagen hydrogels when compared to 2.5 mg/mL collagen when treated with IS, though there was not a significant difference comparing IS to untreated HUVECs (**Figure 4C**). E-selectin expression levels were notably reduced with IS treatment on 2.5 mg/mL hydrogels and were significantly lower on 6 mg/mL hydrogels compared to 2.5 mg/mL (**Figure 4D**). Based on the western blot analysis, we next directly compared monocyte adhesion as a function of collagen density and IS treatment and found that monocyte adhesion followed a similar trend to the ICAM-1 expression levels (**Figure 4E-F**). These results suggest that the uremic environment increases monocyte adhesion in a synergistic manner with ECM stiffness and is driven by ICAM-1 expression by endothelial cells.

**Figure 4.**
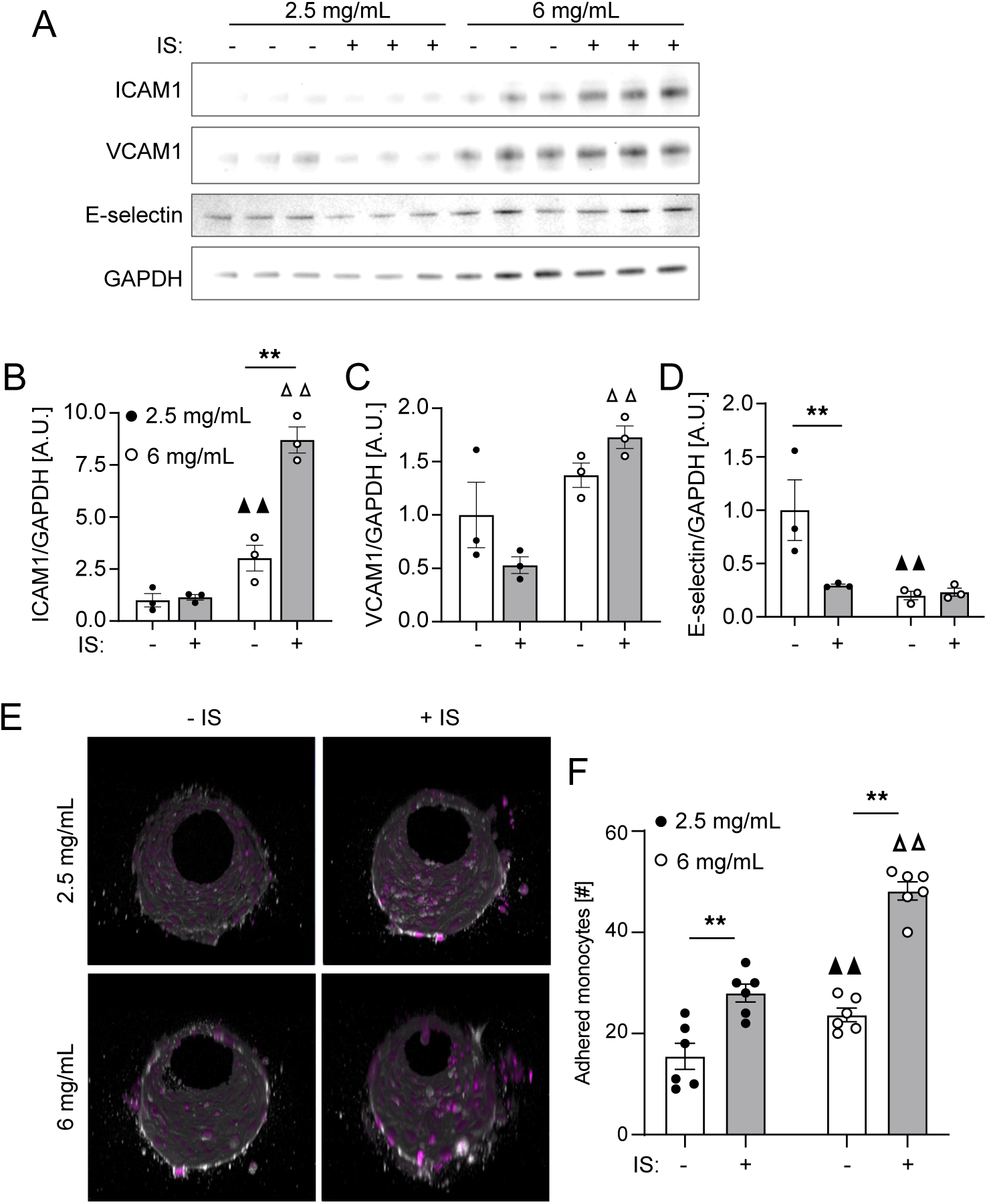
- Effect of uremic microenvironment on expression of adhesion receptors by endothelial cells. **A)** Western blot and **B-D)** quantification of Western blot bands for adhesion molecules in HUVECs treated with and without indoxyl sulfate (2.5 µg/ml) on 2.5 mg/ml and 6 mg/ml collagen hydrogels (three wells per condition; ▲▲ p < 0.01 versus untreated HUVECs on 2.5 mg/ml collagen, Δ Δ p < 0.01 versus Indoxyl sulfate (2.5 µg/ml) treated HUVECs on 2.5 mg/ml collagen, **p < 0.01, two-way ANOVA with a Bonferroni post-hoc test). **E)** 3D reconstructions of confocal micrographs of microvessel stained for DAPI (magenta) and VE-cadherin (white) and THP-1 monocytes stained for DAPI (magenta) and **F)** quantification of adhered monocytes to HUVEC microvessel wall with and without indoxyl sulfate treatment in 2.5 mg/ml and 6 mg/ml collagen hydrogels for 0 Pa pressure condition (three microvessel per condition; ▲▲ p < 0.01 versus untreated microvessels in 2.5 mg/ml collagen, Δ Δ p < 0.01 versus indoxyl sulfate (2.5 µg/ml) treated microvessels in 2.5 mg/ml collagen, **p < 0.01, two-way ANOVA with a Bonferroni post-hoc test). All plots, mean ± SEM. Scale bar 50 µm.

### 2.5 Independent effects of IS on monocytes and endothelial cells

While ICAM-1 expression levels could explain the differences in monocyte adhesion, the differences in monocyte migration distance after extravasation suggest an independent effect of IS on monocyte migration. To dissect the effects of IS on monocytes and the endothelium, we first treated monocytes with IS for 24 hrs then introduced treated monocytes into untreated microvessels, and conversely, we pretreated microvessels with IS for 24 hrs then introduced untreated monocytes (**Figure 5A**). For both treatment conditions, monocytes attached and transmigrated across the microvascular wall into the underlying matrix (**Figure 5A**). Interestingly, treatment of the microvessel with IS significantly increased the number of monocytes that attached to the microvessel wall when compared to IS-treated monocytes for both ECM densities (**Figure 5B-C**). Moreover, we noted a significant increase in monocyte extravasation in 6 mg/mL hydrogels as compared to 2.5 mg/mL for both treatment conditions (**Figure 5D**). Treatment of the microvessel with IS also resulted in significantly increased extravasation as compared to treatment of monocytes (**Figure 5E**). Interestingly, treatment of monocytes with IS resulted in significantly greater migration distances for both collagen densities as compared to treatment of the microvessel (**Figure 5F**). Moreover, the migration distance was greater in 2.5 mg/mL collagen hydrogels compared to 6 mg/mL for both treatment conditions (**Figure 5F**). Collectively, these results suggest that monocyte adhesion and migration are regulated by independent effects on the endothelium and monocytes, respectively.

**Figure 5.**
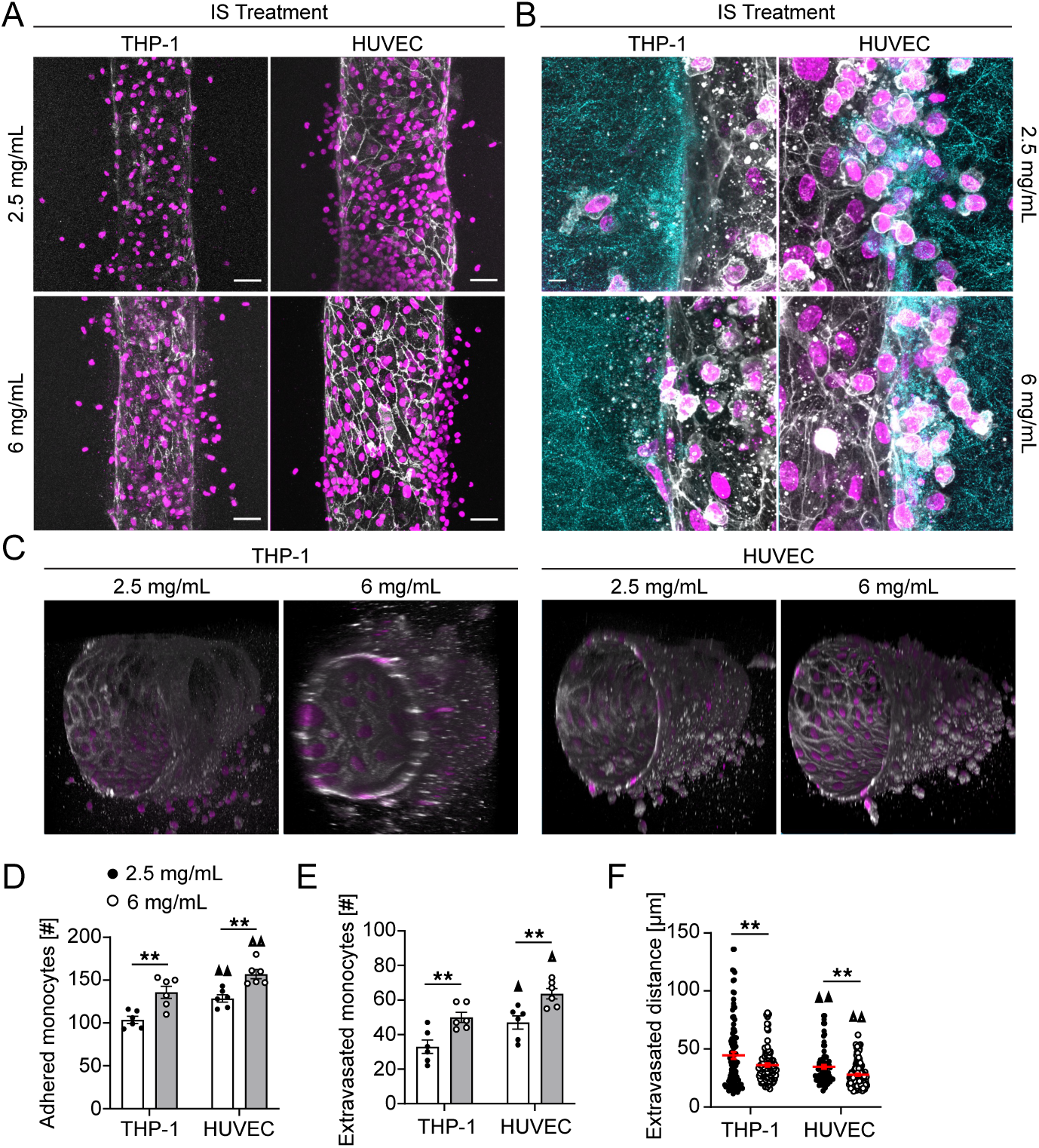
- Effects of indoxyl sulfate pretreatment on THP-1 monocytes and microvessels. **A)** Maximum intensity projections of confocal micrographs of HUVECs stained with DAPI (magenta) and VE-cadherin (white) and THP-1 monocytes stained with DAPI (magenta, scale bar 50 µm), **B)** Magnified micrographs of THP-1 monocytes that extravasated into the interstitial space (Scale bar 10 µm), **C)** 3D reconstructions of confocal micrographs of microvessel stained for DAPI (magenta) and VE-cadherin (white) and THP-1 monocytes stained for DAPI (magenta), **D**) adhered monocytes to endothelium, **E**) extravasated monocytes across endothelium, and **F**) monocyte extravasation distance for THP-1 monocytes pretreated with indoxyl sulfate (2.5 µg/ml) and later added to an otherwise normal HUVEC microvessel and indoxyl sulfate (2.5 µg/ml) treated HUVEC microvessel containing otherwise normal THP-1 monocyte in 2.5 mg/ml and 6 mg/ml collagen matrices for 20 Pa pressure condition (three microvessel per condition; ▲ p < 0.05, ▲▲ p < 0.01 versus Indoxyl sulfate pretreated monocyte in 2.5 mg/ml collagen, Δ p < 0.05, Δ Δ p < 0.01 versus Indoxyl sulfate pretreated monocyte in 6 mg/ml collagen, **p < 0.01, two-way ANOVA with a Bonferroni post-hoc test. All plots, mean ± SEM).

### 2.6 Effects of ECM property and uremic environment on podosome formation and phagocytosis capability

Under stimulation with p-cresyl and IS, monocytes exhibit increased podosome formation and enhanced transmigration through an ILK/AKT signaling pathway-dependent mechanism, which could contribute to vascular damage [55]. To assess whether extracellular matrix properties influenced podosome formation and if IS treatment affected this response, we cultured THP-1 monocytes embedded within 2.5 and 6 mg/mL collagen hydrogels (**Figure 6A**). The percentage of monocytes with podosomes was significantly higher in the 6 mg/ml hydrogel compared to the 2.5 mg/ml hydrogel (**Figure 6B**). Additionally, treatment with IS significantly increased the percentage of monocytes with podosomes at both collagen densities compared to untreated cells **(Figure 6A-B)**. We next examined how substrate properties and uremic environment influenced phagocytosis by evaluating the uptake of microbeads by THP-1 monocytes (**Figure 6C**), as substrate properties have previously been shown to impact the phagocytic capacity of THP-1- derived macrophages, which in turn determines their pro-inflammatory or anti-inflammatory phenotype [53]. The number of beads internalized per cell was significantly lower in the 6 mg/ml hydrogels (∼25 beads/cell) compared to the 2.5 mg/ml hydrogels (∼32 beads/cell) in untreated conditions, indicating a reduced phagocytic activity and a pro-inflammatory like phenotype in 6 mg/ml hydrogels (**Figure 6D**) [53]. IS stimulation further decreased phagocytosis at both concentrations, with bead uptake dropping from 32 to 25 beads/cell for the 2.5 mg/mL hydrogel and from 25 to 18 beads/cell for the 6 mg/mL hydrogel, suggesting a synergistic effect of extracellular matrix properties and uremic microenvironment on such a response. Furthermore, IS treatment significantly reduced the monocyte spread area compared to the untreated case at both collagen densities (**Figure 6E**). Regardless of IS treatment, THP-1 monocytes in the 6 mg/ml hydrogel had the smallest spread area compared to those in the 2.5 mg/ml hydrogel. Interestingly, the cell spread area in the present study contradicts previously reported findings and warrants further investigation [53,56–58]. Collectively, these results suggest that THP-1 cells adapt their migratory behavior in response to the underlying matrix property and the uremic environment may eventually modulate the phenotypic response of such migratory cells.

**Figure 6.**
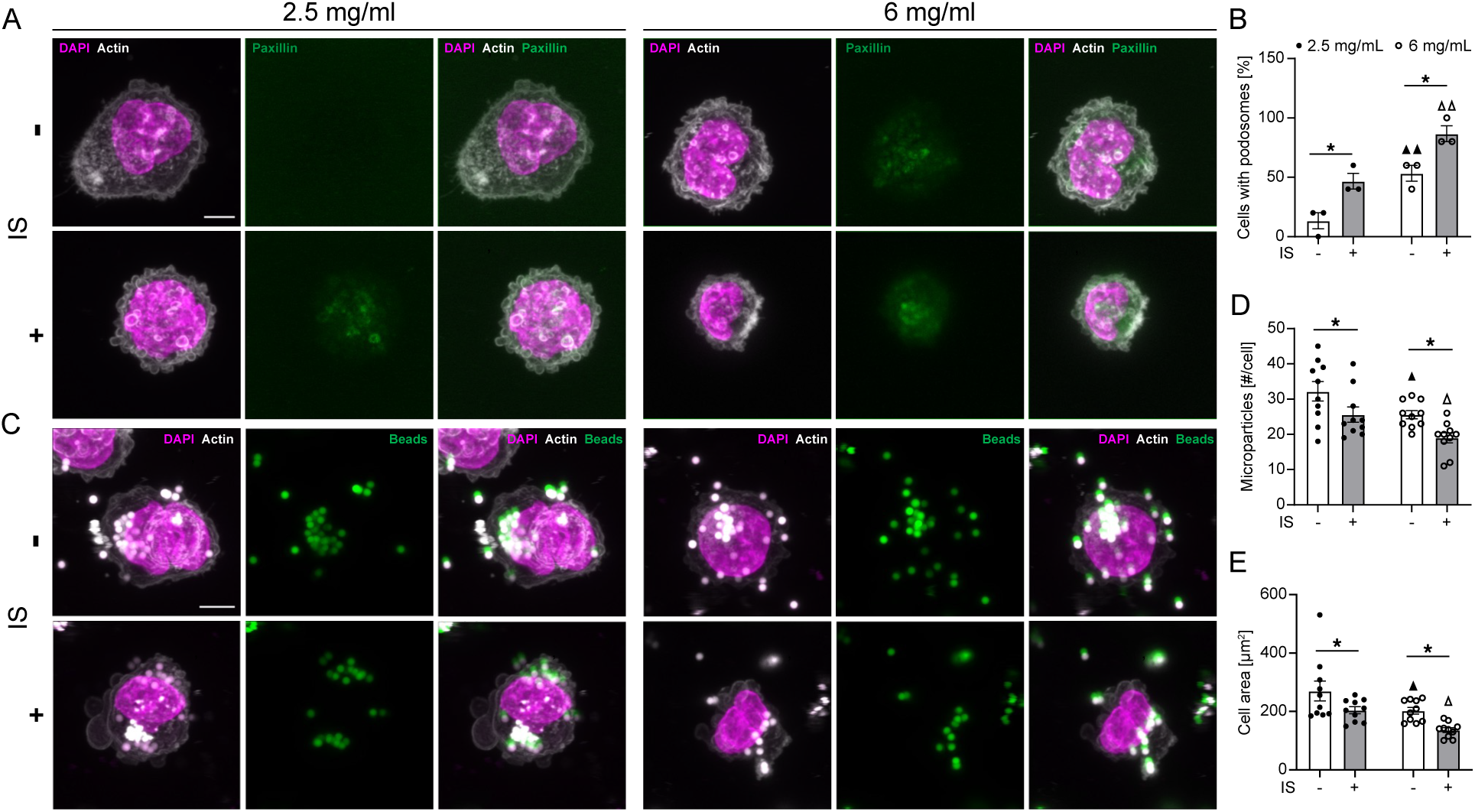
- **Effects of matrix property and uremic environment on monocyte podosome formation and phagocytosis. A**) Podosome formation in THP-1 monocyte stained for DAPI, actin, and paxillin. **B**) Quantification of podosome formation in THP-1. **C**) Maximimum intensity confocal micrographs of THP-1 monocyte phagocytosis of 1µm beads. **D**) Quantification of bead uptake by THP-1 cells. **E**) THP-1 monocyte cell spread area in 2.5mg/ml and 6 mg/ml collagen hydrogels with and without indoxyl sulfate treatment. (three wells per condition; ▲ p < 0.05, ▲▲ p < 0.01 versus untreated case in 2.5 mg/ml collagen, Δ p < 0.05, Δ Δ p < 0.01 versus indoxyl sulfate treatment in 2.5 mg/ml collagen, *p < 0.05, two-way ANOVA with a Bonferroni post-hoc test). All plots, mean ± SEM. Scale bar 5 µm.

## 3. Discussion

Here we present a microfluidic model that simulates monocyte-laden three-dimensional microvessels and demonstrate its capability to mimic the processes of monocyte adhesion, extravasation, and migration through subluminal matrix in a hypertensive uremic setting. Notably, this model addresses limitations of traditional assays by enabling the manipulation of critical physiological factors such as pressure, flow, and the properties of the subluminal extracellular matrix in the presence of the uremic toxin indoxyl sulfate, which is frequently associated with inflammation-related cardiovascular disease in chronic kidney disease. Our previous findings have indicated that the density of the subluminal matrix influences vessel integrity, which is known to impact monocyte extravasation [51]. Additionally, we have also demonstrated a direct correlation between vessel integrity and various stages of CKD [59], collectively suggesting that the uremic environment may independently contribute to inflammation-related CVD within CKD.

In an ex vivo model, it has been demonstrated that high intraluminal pressure alone, without the influence of external hormonal factors, can trigger the expression of chemokines and adhesion molecules, leading to monocyte adhesion to the vascular walls of mouse carotid arteries [60]. Consistent with these results, we found that changes in pressure alone were enough to significantly impact monocyte adhesion and extravasation, regardless of the subluminal matrix condition (**Figure 2**). The uremic environment further intensified the attachment and migration of monocytes across the vascular wall **(Figure 3**), suggesting that elevated luminal pressure may play a pivotal role in triggering processes that lead to CVD and, when combined with circulating uremic toxins, could synergistically enhance inflammation-driven CVD in CKD. Therefore, maintaining optimal luminal pressure may represent a promising therapeutic strategy for preventing CVD related to CKD.

Arterial stiffness is elevated in CKD patients [33–36] and is increasingly recognized as a significant prognostic index and potential therapeutic target in individuals with hypertension [61]. In the present study, denser matrix consistently led to enhanced monocyte adhesion and extravasation (**Figure 2**), and is exacerbated with indoxyl sulfate and elevated pressures(**Figure 3**) suggesting that altered matrix properties, elevated luminal pressure, and uremic environment may work together in recruiting monocytes at endothelium in CKD related CVD. Indoxyl sulfate and p-cresol have been shown to induce THP-1 monocyte endothelial interactions through up-regulation of E-selectin [45] and specific miRNA expression [62]. Notably, here, ICAM-1 was identified as a key driver of THP-1 monocyte adhesion to the endothelium (**Figure 4**). Our data align with previous research showing that IS upregulates ICAM-1 and MCP-1 expression via ROS-induced activation of NF-kappaB in vascular endothelial cells [63]. Taken together, the present results further suggest that indoxyl sulfate may exacerbate CKD-related vascular endothelial inflammation by upregulating ICAM-1 in combination with altered matrix properties and elevated luminal pressures.

In clinical studies, it can be challenging to ascertain whether a specific uremic toxin intrinsically has deleterious effects or simply serves as an inert biomarker for the degree of renal dysfunction. The present microfluidic approach enables us to evaluate the effects of specific uremic compounds on the crosstalk between the major cell types involved in vascular injury: monocytes and endothelial cells. Our findings reveal that monocyte adhesion and migration are influenced by distinct effects of the uremic environment on both the endothelium and the monocytes individually (**Figure 5**). Specifically, our data indicates that exposing endothelial cells to indoxyl sulfate results in significantly greater adhesion and migration of monocytes across the vascular wall compared to monocytes treated with IS alone (**Figure 5**). These in vitro results align with previous in vivo findings, where administration of IS to rats [64] or mice with either normal [65] or compromised [45] renal function led to increased adherence of leukocytes to blood vessel walls and promoted their migration out of the vessels [64].

In the present study, although higher matrix density led to increased extravasation, the monocyte migration distance was consistently longer in the more compliant hydrogel, regardless of the applied luminal pressure and IS treatment (**Figures 2-3 and 5**), suggesting differing influences of matrix properties on monocyte extravasation and migration. The extravasation of monocytes in the presence of uremic toxins like p-cresol and IS has been tied to the formation of podosomes - temporary adhesive structures that break down the extracellular matrix and facilitate invasive migration of THP-1 monocytes into subluminal matrix [55]. Furthermore, matrix stiffness has been shown to affect the migration patterns of THP-1 cells; cells on softer (11 kPa) and medium (88 kPa) stiffness hydrogels exhibit fast amoeboid migration that is Rho-associated kinase (ROCK)- dependent and podosome-independent, while on stiffer (323 kPa) gels, cells display a slower mesenchymal migration mode that is podosome-dependent and ROCK-independent [53]. The present study reinforces these findings with the observation that higher density ECM fosters the formation of podosomes (**Figure 6**), resulting in shorter migration distances in 6 mg/mL hydrogels compared 2.5 mg/mL hydrogels. Moreover, matrix stiffness has been found to modulate macrophage phenotypes, with stiffer matrix promoting pro-inflammatory M1 phenotype and softer matrix promoting anti-inflammatory M2 phenotype [53, 57, 66]. In the present study, THP-1 cells embedded in denser hydrogels exhibited diminished phagocytic abilities, indicating a pro-inflammatory M1-like macrophage phenotype. In contrast, cells embedded in lower collagen concentration displayed enhanced capacity for phagocytosis, reflecting an anti-inflammatory M2- like macrophage phenotype. **(Figure 6).** Nevertheless, exposure to the uremic environment decreases phagocytic capacity and prompts cells towards a pro-inflammatory like phenotype, irrespective of subluminal matrix density, suggesting that the uremic environment may ultimately surpass the modulatory effects of matrix properties on myeloid cell phenotype.

Currently, a challenge in the clinical management of CKD is balancing cardiovascular and renal protection. Notably, the kidneys may play a critical "cause and effect" role, as patients with CKD have an elevated cardiovascular risk, and hypertension is a primary risk factor for cardiovascular events and stroke, which can further exacerbate renal impairment. The microfluidic model introduced in this study, holds promise for identifying optimal blood pressure targets for hypertensive patients with cardiovascular risks and clarifying the delicate balance between reducing cardiovascular events and safeguarding renal function.

## 4. Conclusion

In summary, we developed a 3D microfluidic model to study the combined effects of hypertensive uremic conditions on inflammatory processes. We demonstrated that uremic toxins exacerbate monocyte adhesion and extravasation across the vascular wall in a hypertensive environment, and that the matrix surrounding the endothelium significantly influences monocyte migration and phenotype. This model provides a valuable tool for exploring the various mechanisms involved in monocyte behavior in CKD-related CVD.

## 5. Experimental Section Microfluidic device fabrication

Microfluidic devices were fabricated from polydimethylsiloxane as described previously [67]. Briefly, Sylgard 184 (Dow Chemical Company, Midland, MI, US) was mixed in a 1:10 crosslinker to base ratio. The mixture was placed in a vacuum chamber for 1 hour then poured onto the microfabricated device molds, and cured overnight at 60°C. The following day, PDMS devices were removed from the mold using a razor and sharp tweezers. The PDMS was cut using biopsy punches (ThermoFisher,Waltham, MA, US) for media and gel ports. Devices and glass coverslips (24 x 30mm, ThermoFisher) were cleaned with clear tape and isopropanol prior to exposure to air plasma for 30 seconds. PDMS devices were then bonded to glass coverslips and incubated at 100°C for at least 20 minutes. The interior of each device was treated with 2 mg/mL dopamine hydrochloride (Sigma Aldrich, St. Louis, MO, US) in Tris-HCl buffer (pH 8.5, bioWORLD, Dublin, OH) for 1 hr at room temperature to promote collagen adhesion to the PDMS surface, then washed with DI-H_2_O. An 80 μm needle buffer (5.5” x 48” smart vinyl, Cricut, South Jordan, UT) was placed in each of the media ports, and the devices were sterilized using ultraviolet (UV) light for 20 minutes followed by autoclaving. Acupuncture needles (160 μm diameter, Dongbang Medical, Korea) were soaked in 70% ethanol for at least 20 minutes for sterilization. The needles were then wiped to remove excess liquid and coated with 0.01% BSA (Sigma-Aldrich) for 30 minutes at room temperature prior to insertion into the devices.

### Hydrogel synthesis

Collagen hydrogels were fabricated from rat tail derived collagen type I (Collagen I, high concentration, Corning, NY, US) on ice as described previously [51, 68]. Reconstitution buffer was made by dissolving 1.2 g of NaHCO_3_ and 4.8 g of 4-(2-hydroxyethyl)-1- piperazineethanesulfonic acid (HEPES, Sigma-Aldrich) in 50 mL of DI-H_2_O. Similarly, 10x Dulbecco’s Modified Eagle Medium (10x DMEM, Sigma-Aldrich) was dissolved in DI-H_2_O. RB and 10x DMEM were added to a 1.5 mL microcentrifuge tube on ice at 10% of the volume of stock collagen I, and 1x PBS was added to achieve the desired final concentration of collagen, and the solution was buffered to pH 8.0 with 1 N NaOH. For visualizing the ECM, stocks of AlexaFluor 647 (AF647)-conjugated collagen type I were prepared as described previously [68] and used at dilutions of 10% of labeled collagen in total collagen I volume. Pre-gel solutions were then injected into microfluidic devices and incubated at 37 °C for 30 minutes before the addition of cell culture medium to each media port for hydration. Needles were left in the devices for at least four hours before removal to ensure sufficient crosslinking of the hydrogel, and vacuum grease was used to seal the needle guide on the device, and devices were incubated overnight in endothelial growth medium (EGM-2, Lonza, Basel, Switzerland) prior to seeding cells.

### Cell culture and seeding

Human umbilical vein endothelial cells (HUVECs, Lonza) were cultured in EGM-2 media from passages 2-9 and passaged at 80-90% confluency. To seed devices, HUVECs were incubated with 0.05% trypsin-EDTA (ThermoFisher) for 1 minute at 37°C, then mechanically perturbed until the cells were lifted as determined by phase contrast microscopy. After cell lifting, the trypsin was inactivated by the addition of growth medium and the cell suspension was centrifuged at 1000 RPM for 5 minutes. Following trypsin inactivation and centrifugation, cells were resuspended to a concentration of 1.8 × 10^6^ cells/mL in EGM-2. Before seeding the cell in microfluidic devices, gel-filling pipette tips were inserted in the hydrogel ports and gently aspirated 3x to clear any media and hydrogel in the ports and to establish a pressure gradient across the vessel wall to increase seeding efficiency. Media was removed from the inlet and outlet media ports and 60 μL of cell suspension was introduced in one of the media ports, and 50 μL cell suspension was introduced in the other port of the same channel to create a slight pressure differential and induce cells to flow into the microchannel. The devices were inverted every minute for 8 min, then incubated upside down in a humidified incubator for 6–8 min followed by inverting and incubating for another 6–8 min in the incubator. Devices were then moved to a laboratory rocker and allowed to oscillate at 5 cycles per min for 2–4 hr before culture media was replaced with fresh EGM-2 media. The devices were kept on the rocker overnight and checked optically the following day. Devices were re-seeded following the process above if the monolayer did not coat the entire vessel.

THP-1 (ATCC® TIB202™, Manassas, VA, USA) is a commercially available monocyte cell line derived from peripheral blood. THP-1 cells were cultured in suspension using RPMI-1640 medium containing L-glutamine, 10% fetal bovine serum (FBS), and ampicillin/streptomycin (RPMI-1640, Gibco, Gaithersburg, MD, USA), and were used between passage numbers 0 and 7.

### Cell fixation and staining

Prior to fixation, devices were washed with PBS++ (PBS with 0.5 mM MgCl_2_ and 1mM CaCl_2_) then incubated with 4% w/v paraformaldehyde in PBS++ for 15 minutes at 37°C and subsequently washed 3x with PBS++. Cells were permeabilized by adding 0.3% Triton X-100 (ThermoFisher) in PBS for 10 minutes on rocker placed in an incubator maintained at 37 °C followed by a wash with PBS++ and later blocked by adding 2% bovine serum albumin (BSA) in PBS for at least 1h at RT on rocker. The devices were then washed and incubated with mouse anti-VE cadherin (Santa Cruz Biotechnology, Dallas, TX, US) at a 1:200 dilution (v/v) in PBS++ overnight at 4°C on rocker. Devices were next washed 3x with PBS++ and incubated with Alexa Fluor 647-conjugated goat anti-mouse IgG (ThermoFisher) and Alexa Flour 488-conjugated phalloidin (ThermoFisher) at 1:1000 (v/v) and 1:200 (v/v) dilutions, respectively, in PBS++ and incubated for 1 h at room temperature on rocker. The devices were then washed with PBS++ and later incubated with DAPI (ThermoFisher) at a 1:1000 dilution in PBS++ for 10 minutes at RT on rocker followed by a PBS++ wash. Imaging was performed on a laser scanning confocal microscope with a 10x/0.4 NA air objective or a 30x/1.05 NA silicone oil objective (FV3000, Olympus, Tokyo, Japan). ImajeJ was used to synthesize the maximum intensity projections and create 3D reconstruction from Z-stack series obtained with the laser scanning confocal microscopy.

### Adhesion and Extravasation assays

For the adhesion and extravasation assays, a solution of 1.5 × 10^6^ cells/mL of THP-1 monocytes was resuspended in EGM-2 media and injected into bare microchannels or HUVEC microvessels formed in 2.5 and 6 mg/mL collagen hydrogels. For test conditions involving indoxyl sulfate treatment, THP-1 monocytes were resuspended in EGM-2 spiked with 2.5 µg/mL of indoxyl sulfate and subsequently introduced into the devices. Devices were incubated at 37 °C for 24 h after the addition of IS prior to analysis.

### Western Blot

Cells were rinsed once with ice cold PBS and lysed with radioimmunoprecipitation assay buffer (RIPA buffer, Thermo Fisher Scientific) supplemented with 1X HALT protease and phosphatase inhibitor (Thermo Fisher Scientific). Lysates were homogenized by pulse-vortexing and clarified by centrifugation at 14,000 x g for 5 min at 4 °C. The concentration of the clarified lysates was measured using Pierce BCA Protein Assay (Thermo Fisher Scientific). Lysates were then reduced with NuPAGE LDS reducing agent and dithiothreitol, followed by denaturation for 5 minutes at 100 °C. The denatured proteins were separated on NuPAGE 4 to 12% bis-tris gradient gels (Thermo Fisher Scientific) and transferred to iBlot2 PVDF ministack membranes. Membranes were blocked with SuperBlock Blocking Buffer (Thermo Fisher Scientific) for an hour with rocking at room temperature and then incubated overnight with the following primary antibodies with rocking at 4 °C: anti-VCAM-1(1:1000, Cell Signaling Technology, #13662), anti-E-selectin (1 µg/mL, Novus Biologicals, BBA16), anti-ICAM-1 (1:1000, Cell Signaling Technology, #4915), and anti-glyceraldehyde-3-phosphate dehydrogenase (1:5000, Cell Signaling Technology, #2118) diluted in blocking buffer. After overnight incubation, membranes were washed three times with TBS-T and subsequently incubated with horseradish peroxidase conjugated antibodies (1:10,000, v/v, diluted in 5% milk (w/v)) for 1 hour at room temperature. Membranes were washed three times over 15 minutes with TBS-T and developed using chemiluminescence with either SuperSignal West Femto Maximum Sensitivity Substrate reagent (Thermo Fisher Scientific) or Clarity Western ECL reagent (BioRad). Western blot images were quantified using ImageJ.

### Phagocytosis of beads and podosome formation on monocytes embedded in collagen

A suspension of THP-1 cells at a concentration of 5×10^6^ cells/mL was prepared in PBS. Collagen hydrogels with concentrations of 2.5 mg/mL and 6 mg/mL were synthesized as previously described. To achieve the final desired collagen concentration, the THP-1 cell suspension was substituted for 1x PBS. Yellow-green amine-modified beads (1 micron diameter, FluoSpheres, Invitrogen), were sterilized by incubating at 70°C for 24 hours, followed by two washes with 0.1 M sodium hydroxide (NaOH) to remove potential endotoxin contaminants. The sterilized beads were then diluted in fresh THP-1 media, either with or without IS, to a final concentration of 50 beads per cell. After 24 hours of incubation with the beads, the cells were washed twice with PBS++, fixed, and stained with Phalloidin and DAPI to visualize the cytoskeleton and nucleus. Confocal imaging was performed using a laser scanning confocal microscope with 60x/1.4 NA oil objective (FV3000, Olympus, Tokyo, Japan). Images were analyzed using ImageJ, with bead counts manually performed using the ‘Cell Counter’ plugin.

To analyze podosome formation, a permeability-fixation solution was prepared consisting of 1% w/v paraformaldehyde in PBS++ and 0.05% Triton X-100 in PBS++. THP-1 cells embedded in collagen hydrogels at concentrations of 2.5 mg/mL and 6 mg/mL were washed 3X with PBS++ and then incubated with the permeability-fixation solution for 5 minutes at 37 °C. After this, the solution was replaced with 4% w/v paraformaldehyde in PBS++ and incubated for 45 minutes at 37 °C, followed by 3X washes with PBS++. The cells were then permeabilized by incubating with 0.1% Triton X-100 for 45 minutes at 37 °C. After permeabilization, the cells were incubated with 1% BSA in PBS for at least 1 hr at RT. To identify podosomes, the cells were washed and incubated overnight at 4°C with anti-mouse paxillin (Clone 117, BD Biosciences) at a 1:200 dilution (v/v) in PBS++. The cells were then washed 3X with PBS++ and incubated for 2 hrs at RT with Alexa Fluor 647-conjugated goat anti-mouse IgG (ThermoFisher) and Alexa Fluor 488- conjugated phalloidin (ThermoFisher) at 1:500 (v/v) and 1:200 (v/v) dilutions, respectively.

Afterward, the cells were washed with PBS++ and incubated for 30 minutes at RT with DAPI (ThermoFisher) at a 1:1000 dilution in PBS++. The cells were then washed again with PBS++. Maximum intensity projections were generated in ImageJ from Z-stack series obtained using laser-scanning confocal microscopy (FV3000, Olympus, Tokyo, Japan) with a 60x/1.4 NA oil objective.

### Statistical Analysis

All data are presented as mean ± standard error of the mean (SEM) unless otherwise specified in the Figure caption. Statistical significance was evaluated using student’s t-test or two-way analysis of variance (ANOVA), followed by Bonferroni test, where appropriate. All statistical tests and plots were generated using Prism (GraphPad v10.1.2). Significance was considered for p < 0.05 and indicated in Figure captions. The number of microvessels tested for significance are indicated in Figure captions.

## Supporting information

Supplementary Methods and Results

## Acknowledgements

This work was supported by the National Institutes of Health (R35GM142944), the American Heart Association (CDA857738). M.R. acknowledge financial support of Institute for Convergent Science, UNC Chapel Hill. W.Y.A. acknowledge grants from the Lymphatic Malformation Institute and the CLOVES Syndrome Community. S.A.H., and E.L.D. acknowledge financial support of the National Institutes of Health through the Integrative Vascular Biology Training Program (T32HL69768 to E.L.D. and S.A.H.), the UNC Kidney Center T32 (T32DK007750 to S.A.H.), and a Ruth L. Kirchstein predoctoral individual fellowship (F31HL162462 to E.L.D.). Microfabrication was performed in the Chapel Hill Analytical and Nanofabrication Laboratory, CHANL, a member of the North Carolina Research Triangle Nanotechnology Network, RTNN, which was supported by the National Science Foundation (ECCS-2025064), as part of the National Nanotechnology Coordinated Infrastructure, NNCI.

## Notes

### Competing Interest Statement

The authors have declared no competing interest.

