## Supplementary Methods and Results for "Microphysiological uremia model reveals biophysical potentiators of vascular dysfunction"

### SUPPLEMENTAL METHODS

#### Quantifying hydrogel hydraulic conductivity

To determine the hydraulic conductivity from the height of the fluid column as a function of time (**Fig. 1**), a differential equation resulting from combining Darcy's Law (Eq. 1) with hydrostatic pressure (Eq. 2) and conservation of mass (Eq. 3):

$$u = \frac{Q(t)}{A_{\text{hydrogel}}} = \frac{K \cdot \Delta p(t)}{L} \qquad (1)$$

$$\Delta p(t) = \rho g\big(\Delta h(t)\big) \qquad (2)$$

$$\Delta h(t) = \Delta h_0 - \frac{1}{A_{\text{reservoir}}} \int_0^t Q\,(t)dt \qquad (3)$$

Where $u$ is the fluid velocity magnitude through the porous media, $Q$ is the volumetric flow rate, $A_{hydrogel}$ is the cross sectional area of the hydrogel, $K$ is the hydraulic conductivity, Δp is the pressure drop from the inlet to the outlet, $\rho$ is the density of water, $g$ is the acceleration due to gravity, Δh is the fluid column height difference between the inlet and the outlet, and $A_{reservoir}$ is the total cross sectional area of the inlet reservoirs. Combining these equations yields a differential equation, the solution to which is given by:

$$\Delta h(t) = \Delta h_0 \cdot exp^{-c \cdot t} \qquad (4)$$

Where $c$ is a combination of parameters:

$$c = \frac{\rho \cdot g \cdot K \cdot A_{\text{hydrogel}}}{L \cdot A_{\text{reservoir}}} \qquad (5)$$

To determine the hydraulic conductivity, Eq. 4 was fit to the height difference as a function of time (**Fig. 1**), and $K$ was determined with known values for the constants.

#### Determining microvessel permeability with luminal pressure

To quantitatively determine whether vessel permeability varied as a function of transmural pressure gradients that could drive flux via convection in addition to diffusion, we seeded the pressure source channel with HUVECs and cultured devices overnight on a laboratory rocker. On the following day, 70kDa FITC dextran was added to the devices, which were then returned to the rocker overnight to flood the ECM with a uniform concentration of dextran. On day 3, a cocktail of 70kDa TR and FITC dextran was administered to the seeded channel via reservoirs to apply luminal pressures of 0, 20, and 200 Pa (**Fig. S4**). Transmural dextran flux was determined using confocal microscopy, with reflectance of 488 nm light used to image the collagen hydrogel in a label-free manner and FRAP of the FITC channel used to measure transmural velocity magnitude (**Fig. S4**). While we found that transmural flux significantly increased with luminal pressure for microvessels embedded in 2.5 and 6 mg/mL collagen gels, we did not observe a significant difference in flux between the two collagen densities at any applied pressure. Similarly, we observed increased transmural fluid velocity at each applied pressure, but no significant difference between collagen densities (**Fig. S4**). Interestingly, we found a strong correlation between measured transmural fluid velocity magnitude and total dextran flux that was independent of collagen density (**Fig. S4**).

### SUPPLEMENTAL RESULTS

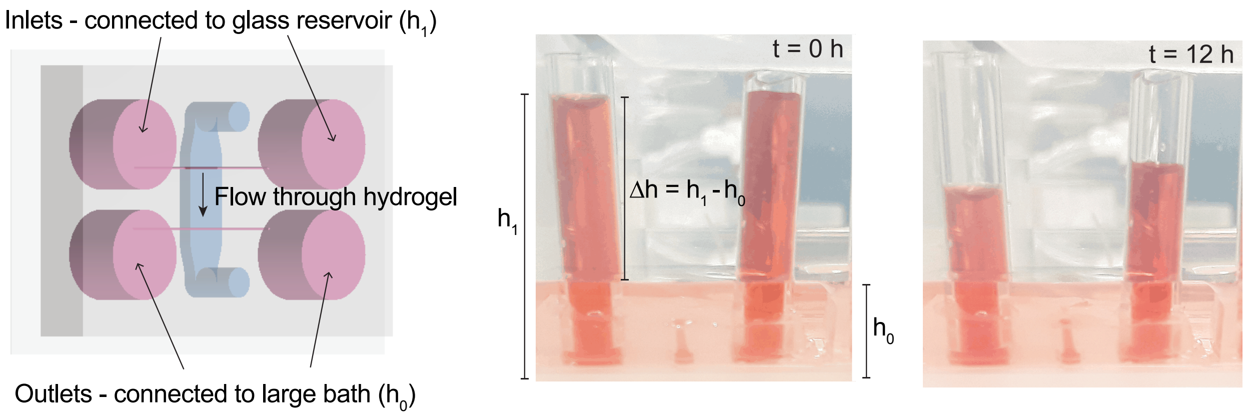


**Figure S1.** Method for determining bulk hydraulic conductivity. Microfluidic devices containing 2 parallel 160 μm diameter channels separated by 4 mm were connected to 6 mm OD glass tubing at the inlet and bathed in PBS at the outlet. The difference in height of the fluid column between the inlet and outlet was measured as a function of time, and the resulting height vs. time curves were used to quantify the hydraulic conductivity.

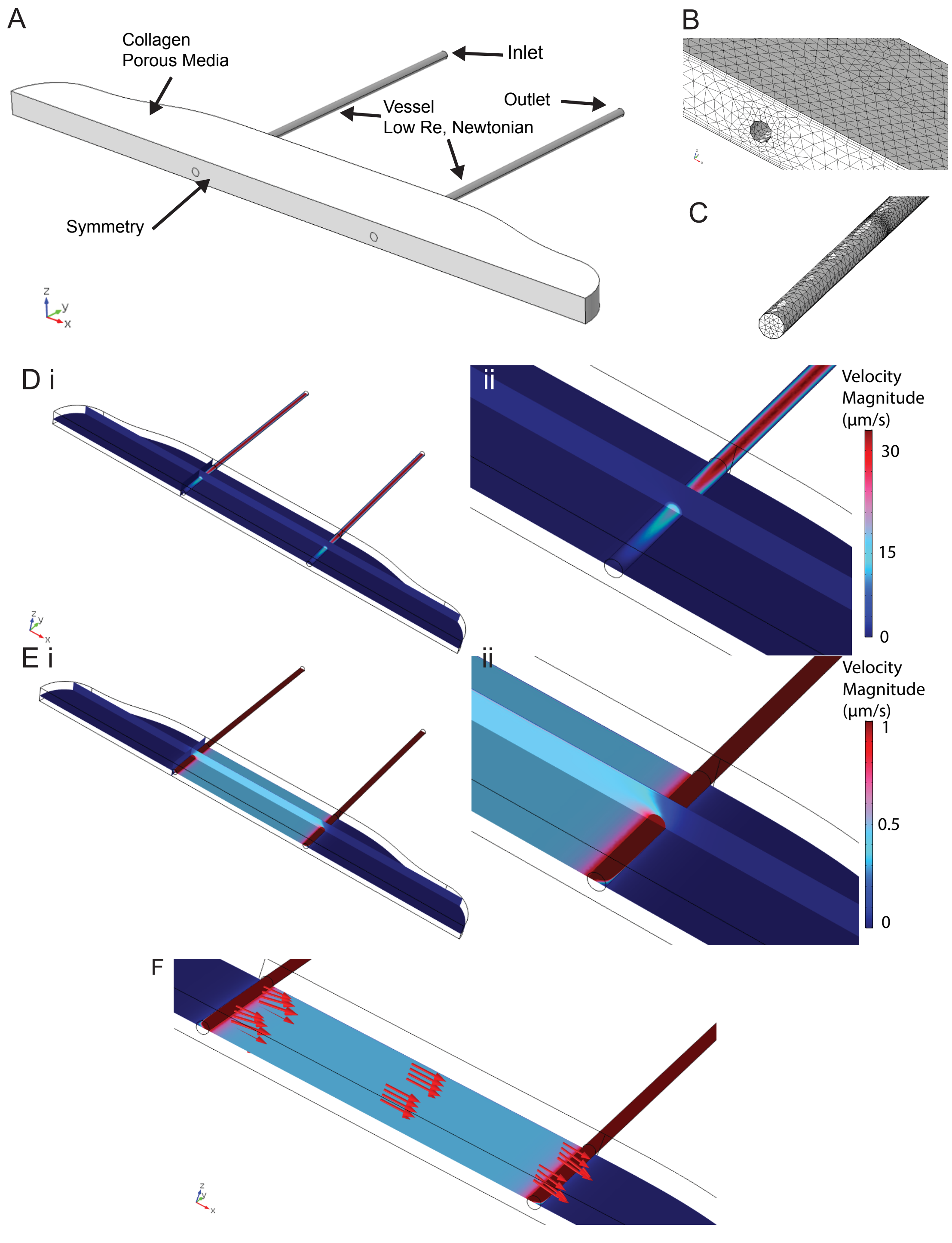


**Figure S2: 3D model of the two-channel device. A)** Overview of vessel geometry: the central region is modeled as a porous medium and the intersecting channels as open low Re domains. **B-C)** Mesh of the gel region and of the channel. The intersection of the channel and porous medium is examined with a higher density of meshes. **(D)** Full range of velocity magnitudes across device at x-y and x-z cross sections. (i) Overview of transport across device. (ii) View of velocity magnitude along one channel. **(E)** Velocity magnitudes across device with adjusted visual range. (i) Overview of transport across device. (ii) View of velocity magnitude at 13-open channel interface. **(F)** Arrow volume depicting velocity vectors through the collagen hydrogel.

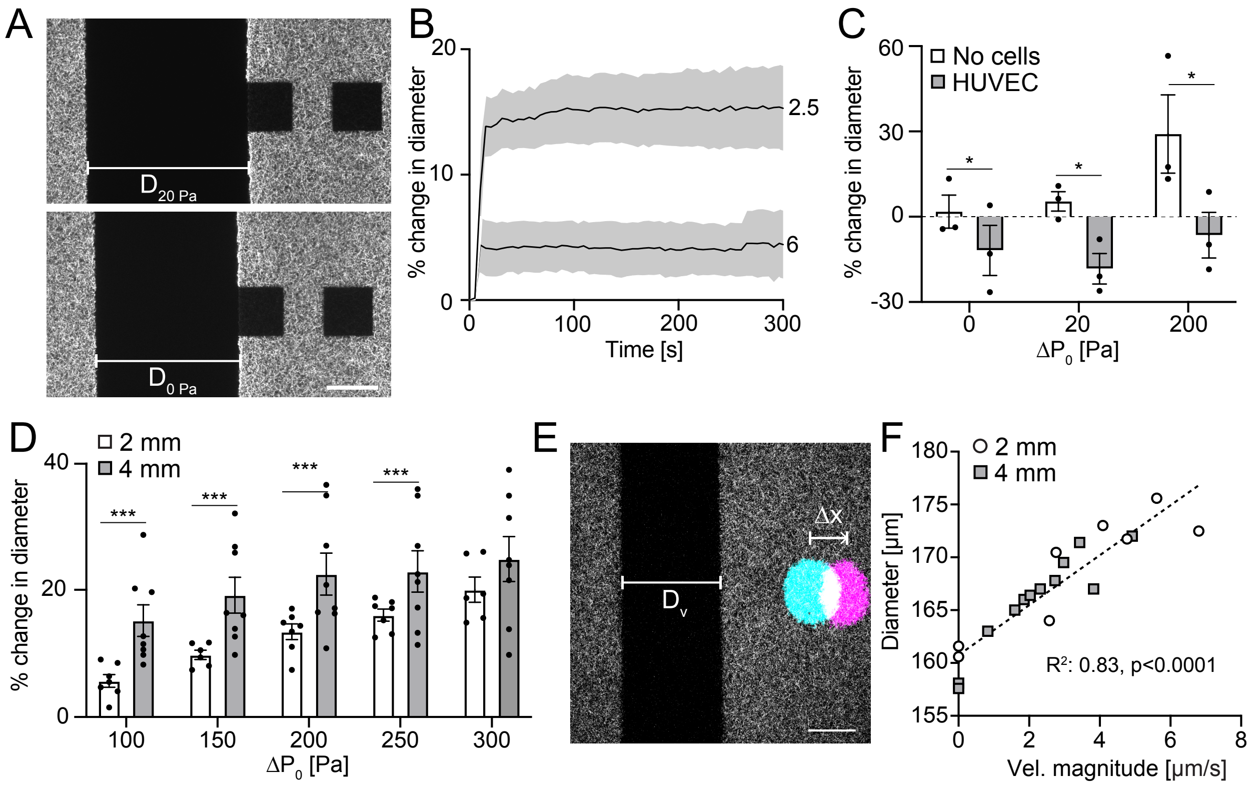


**Figure S3. Transmural pressure gradients modulate vessel strain. A)** Confocal micrographs of fluorescent collagen during and after loading with 20 Pa pressure in the vessel lumen. Parallel, second channel was maintained at 0 Pa. Black squares are photobleached fiduciary markers for aligning images before and after loading (scale bar 50 μm). **B)** Change in diameter of lumen after application of 20 Pa of luminal pressure while maintaining parallel channel at 0 Pa for 2.5 and 6 mg/mL collagen hydrogels (black line indicates mean, grey corridors are +/- S.E.M.). **C)** Change in lumen diameter with and without HUVECs seeded in channels formed in 2.5 mg/mL collagen hydrogels. **D**) Change in lumen diameter in single-channel devices with 2 mm or 4 mm outlet ports. **E**) Representative confocal micrograph of fluorescent collagen (grey) and photobleached 70 kDa dextran (blue – inverted bleached spot at t = 0, magenta – inverted bleached spot at t = 20 s, scale bar 50 μm). **F**) Plot of lumen diameter as a function of interstitial fluid velocity magnitude as measured by FRAP (black dots indicate measurement made from independent device, *p<0.05, **p<0.01, ***p<0.001 as determined by ANOVA).

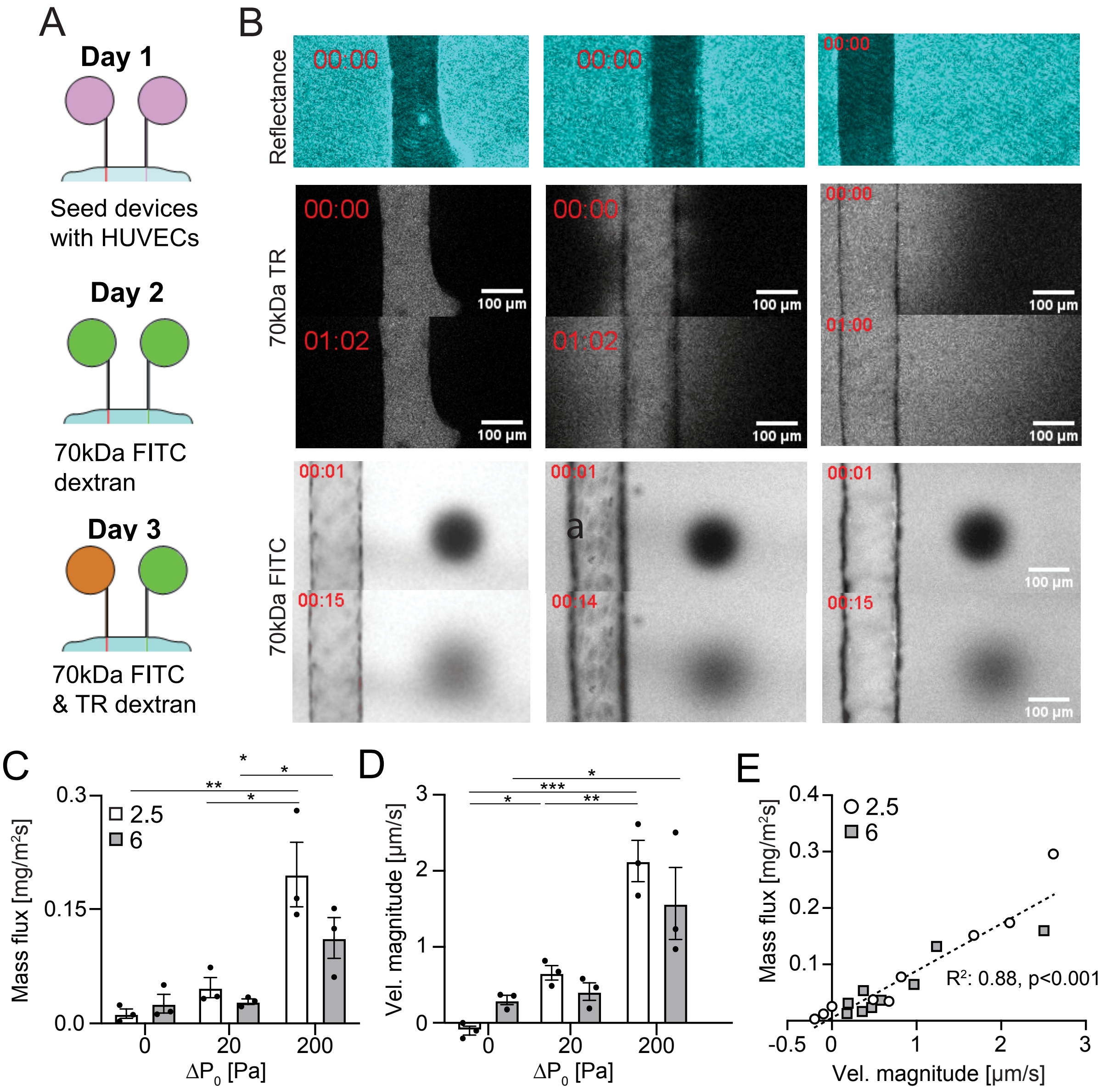


**Figure S4. Assay to measure transmural flux in double channel devices as a function of luminal pressure**. **A**) After seeding devices with HUVECs, media containing 70 kDa FITC dextran was added to all ports in the device to achieve a uniform concentration by the following day at which point a solution of 70 kDa FITC dextran and TR dextran was added to the channel containing cells. **B**) Confocal reflectance imaging was used to visualize the ECM and confocal imaging was used to measure the flux of TR dextran from the lumen into the surrounding ECM. A modified FRAP protocol of FITC dextran was used to quantify fluid velocity magnitude and direction. **C**) Transmural flux of 70 kDa dextran and **D**) fluid velocity magnitude as measured by FRAP across microvessels seeded with HUVECs. **F**) Mass flux as a function of fluid velocity magnitude for devices seeded with HUVECs in 2.5 and 6 mg/mL collagen (data points represent independent devices, bars are mean +/- SEM, *p<0.05, **p<0.01, ***p<0.001 as determined by ANOVA).
